# Self-organized BMP signaling dynamics underlie the development and evolution of tetrapod digit patterns

**DOI:** 10.1101/2023.03.28.534660

**Authors:** Emmanuelle Grall, Christian Feregrino, Sabrina Fischer, Aline De Courten, Tom W. Hiscock, Patrick Tschopp

## Abstract

Repeating patterns of synovial joints are a highly conserved feature of articulated digits, with variations in joint number and location giving rise to a diverse range of digit morphologies and limb functions across the tetrapod clade. During development, joints form iteratively within the growing digit ray, as a population of distal progenitors alternately specifies joint and phalanx cell fates to segment the digit into distinct elements. Whilst numerous molecular pathways have been implicated in this fate choice, it remains unclear how they give rise to a repeating pattern. Here, using single cell RNA-sequencing and spatial gene expression profiling, we investigate the transcriptional dynamics of interphalangeal joint specification *in vivo*. Combined with mathematical modelling, we predict that interactions within the BMP signaling pathway – between the ligand GDF5, the inhibitor NOG, and the intracellular effector pSMAD – result in a self-organizing Turing system that forms periodic joint patterns. Our model is able to recapitulate the spatiotemporal gene expression dynamics observed *in vivo*, as well as phenocopy digit malformations caused by BMP pathway perturbations. By contrasting *in silico* simulations with *in vivo* morphometrics of two morphologically distinct digits, we show how changes in signaling parameters and growth dynamics can result in variations in the size and number of phalanges. Together, our results reveal a self-organizing mechanism that underpins tetrapod digit patterning and its evolvability, and, more broadly, illustrate how Turing systems based on a single molecular pathway may generate complex repetitive patterns in a wide variety of organisms.

## Introduction

Articulated digits with repeating joints are a hallmark of the distal tetrapod limb and considered an essential early adaptation to locomotion on dry land (1–3). Digits are segmented into individual digit bones, termed phalanges, which are connected to each other by synovial joints that facilitate relative motion of adjacent bones and, thus, digit flexion. Thanks to this modular architecture, highly distinct digit patterns have evolved to enable locomotory behaviors as diverse as walking, swimming, flying, or the execution of fine motor skills (4). These patterns are individualized both in terms of number and size of their phalangeal bones and can vary across species as well as within the same limb. For the latter, differences in phalangeal formulas are a manifestation of distinct digit identities along the anteroposterior axis of the distal limb domain, the so-called autopod (4, 5).

The foundation for this diversity in digit morphologies is laid down early during embryonic patterning. Once the three major proximal-distal segments of the tetrapod limb have been defined – the stylopod (upper arm/leg), zeugopod (forearm/lower leg) and autopod (hand/foot) – digit outgrowth is initiated at the distal margin of the autopod, through the specification of a progenitor population at the distal digit tip known as the phalanx forming region (PFR) (6, 7). Molecular and mechanical cues converge to shape this organizing center, with fibroblast growth factors (FGFs) from the overlaying apical ectodermal ridge (AER) defining a distal domain of growth competency (8–10). Cell rearrangements at the PFR result in rod-shaped and elongating digital rays, as progenitors become incorporated distally (9) (Fig.1A). Once digit progenitors leave the signaling environment of the PFR niche, they differentiate into either chondrocytes, the phalanx progenitors, or interzone cells, which mark the site of future interphalangeal joints (11). By alternately differentiating into chondrocyte or interzone cell fates, the digit thus forms a repeating sequence of phalanges and joints. Accordingly, the number and initial size of skeletal elements is defined by the periodicity and duration with which either of the two cell types is sequentially specified, as the digital ray elongates (Fig.1A,B).

**Figure 1.**
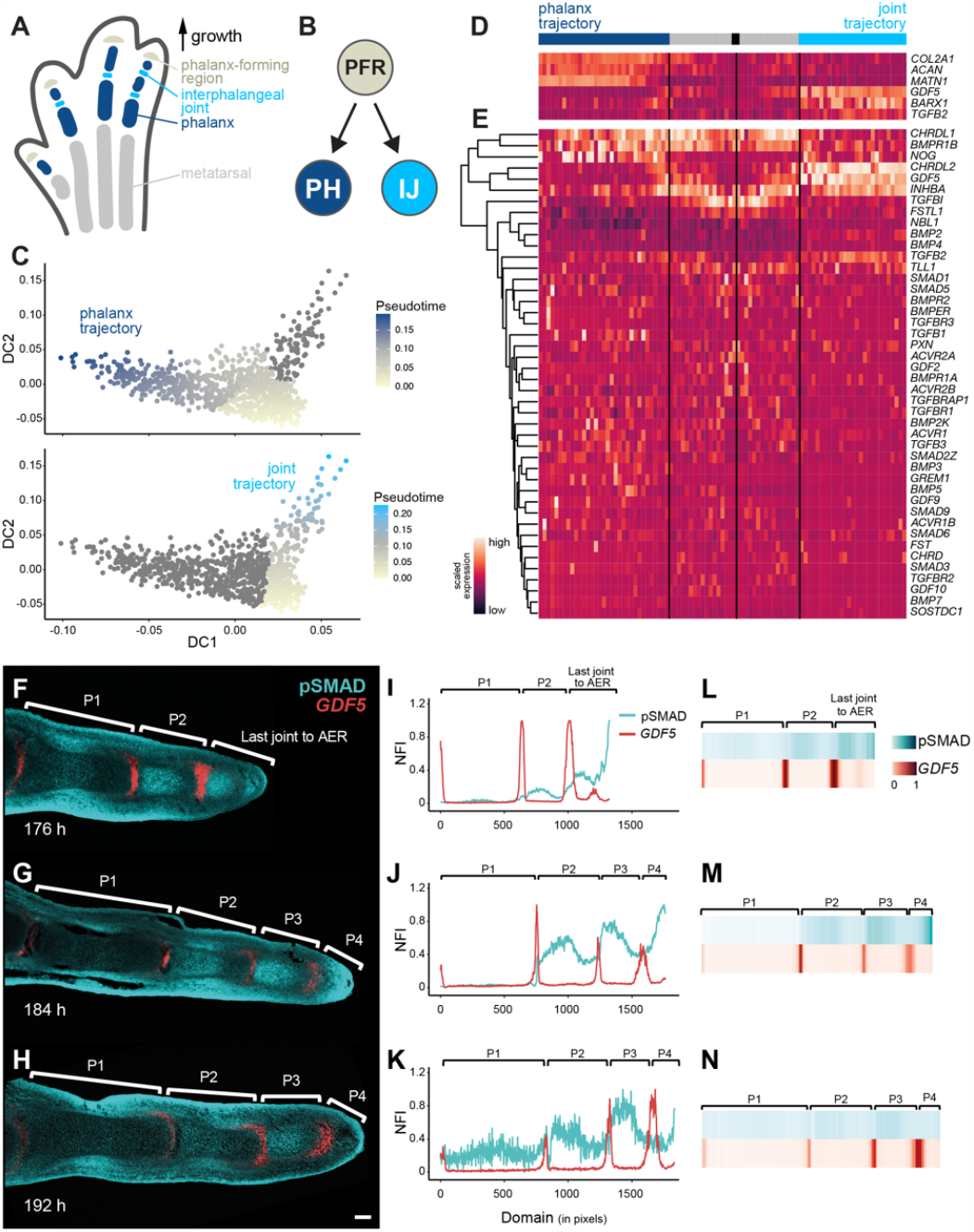
TGF-beta signaling dynamics during the phalanx/joint cell fate decision. (**A-E**) Pseudotime single-cell RNA-seq analyses of the phalanx/joint-progenitor cell fate decision in the chicken hindlimb. (**A**) Schematic of the different elements present and shaping a developing digit at stage HH29. (**B**) A bifurcating cell fate decision splits the phalanx-forming region (PFR) cell population into either phalanx (PH) or interphalangeal joint (IJ) progenitor cells. (**C**) Pseudotime progression along the phalanx and the joint trajectories. (**D, E**) Pseudotime heatmap of marker genes differentially expressed between the phalanx progenitor trajectory (dark blue) and the joint progenitor trajectory (turquoise). The black box corresponds to the starting point of the pseudotime, the grey zone to the part shared by the two trajectories. Scaled gene expression from low (purple) to high (orange). (**E**) Hierarchically clustered pseudotime heatmap of all detected genes of the TGF-beta superfamily, along the phalanx and joint progenitors’ trajectories. (**F-N**) Spatiotemporal *in vivo* data for *GDF5* gene expression and BMP signaling activity in developing digits. (**F-H**) Fluorescent RNA *in situ* hybridization for *GDF5* combined with immunohistochemistry for pSMAD on longitudinal sections of digit III at 176h, 184h and 192h of development. Scale bar = 100 μm. (**I-K**) Plots of normalized fluorescence intensities (NFI) for pSMAD (cyan) and *GDF5* (red) along the proximal-distal axis of digit III. 1 pixel = 1.243 μm. (**L-N**) Heat map visualizations of NFI for pSMAD (cyan) and *GDF5* (red) of digit III.

This repetitive pattern of cell fates is first observable as periodic and regularly spaced stripes of gene expression of joint cell fate markers along the proximal-distal axis of the digit. Since interphalangeal joint number varies widely across tetrapods – ranging from 1 in the human thumb, to over 15 in certain species of whale – it suggests that the locations of these early stripes of expression are not set individually, but rather are the result of an inherently iterative developmental mechanism that places joints at regular spatial intervals along the digit. Previous studies have proposed that a Turing-like mechanism might be responsible for the development of these patterns (12–14). In this scenario, a network of interacting and diffusing biochemical signals forms a reaction-diffusion system that self-organizes into a periodic pattern (15). The core underlying principle is the presence of a long-range negative feedback loop in the system, typically achieved by a rapidly diffusing inhibitor molecule, whose diffusive range sets the characteristic spacing of the pattern. As such, Turing mechanisms allow repetitive structures to form at regular spatial intervals, akin to how joints repeatedly form along the length of the digit. Several experimental observations implicate a Turing system in phalanx-joint patterning. First, insertion of a foil barrier into the developing digit perturbs phalangeal proportions, suggesting that molecular diffusion plays an essential role in patterning (16). Second, experimentally-induced ectopic joints can repress the formation of endogenous joints, suggestive of Turing-like long-range inhibition (17). Third, certain mouse mutants display aberrant joint patterns predicted to be highly specific to Turing mechanisms (12, 18). However, whilst these data suggest that digit patterning might be Turing-like, the specific molecular network responsible for patterning remains poorly defined, and it remains unclear how it could generate a self-organizing reaction-diffusion system.

Genetic studies in humans and mice, as well as experimental embryology in avian models, have identified many genes and molecular pathways involved in digit development (e.g., WNT, IHH, FGF (10, 17, 19–21). In particular, the bone morphogenetic protein (BMP) pathway, comprising members of the TGF-beta superfamily, appears to play an essential role during phalanx-joint specification. One of the earliest known markers of the joint interzone is the BMP ligand *Growth Differentiation Factor 5* (*GDF5*) (22, 23). GDF5 activates BMP-signaling by binding to its preferred receptor, *BMPR1B*, which is expressed within the developing digit ray (6, 24). Moreover, high levels of pSMAD1/5/8, the nuclear effectors of active BMP signaling, and pSMAD2/3, downstream of *Activin*, mark the PFR progenitor population and are essential for digit outgrowth (6, 7, 9). Differences in pSMAD1/5/8 levels at the PFRs of digits in the same autopod mirror distinct digit identities, and precede the emergence of digit-specific morphologies (25). Finally, congenital defects in human digit patterning are frequently associated with mutations in members of the BMP pathway, including *GDF5, BMPR1B*, and *Noggin* (*NOG*), an extracellular BMP inhibitor known to bind GDF5 (26). Based on these observations, it has been hypothesized that regulatory interactions amongst diffusible members of the BMP pathway may play a critical role in generating the periodic phalanx-joint patterns characteristic of tetrapod digits (11, 13).

Here, we combine single-cell transcriptomic data of the PFR and its descendant cell populations – the phalanx and joint progenitors – with quantitative *in situ* measurements to describe the spatiotemporal dynamics of BMP activity during digit segmentation *in vivo*. Building on the observed dynamics, we use mathematical modeling to propose a BMP-based Turing system that can recapitulate patterning *in silico*, as well as phenocopy experimental perturbations affecting its main molecular constituents. Combined with developmental growth and segmentation data from two morphologically distinct digits, we discuss potential cellular and molecular mechanisms underlying the development and evolution of distinct digit patterns in the distal tetrapod limb.

## Results

### Transcriptional signatures and BMP signaling dynamics of the phalanx-joint cell fate decision

To follow the transcriptional dynamics accompanying the specification of PFR progenitors into either phalanx-forming chondrocytes, or interphalangeal joint-inducing interzone cells, we explored a single-cell RNA-sequencing (scRNA-seq) dataset of the chicken foot at Hamburger-Hamilton stage 29 (HH29) (27, 28). Due to the growth dynamics of the chicken digit at that stage, sampling the distal autopod of multiple embryos – with slight developmental heterochronies between them – should capture the transcriptional signatures of the PFR, the phalanx- and joint-forming progenitors, as well as the intervening cell states of this cell fate decision process (Fig.1A,B, (27)). Indeed, based on marker gene expression analyses, we were able to identify three scRNA-seq cell clusters with transcriptional profiles reminiscent of a proliferative distal cell population with an early chondrocyte signature, a maturing chondrocyte population, and cells showing signs of interzone cell fate induction (Fig.S1A). Focusing on these three clusters, we selected variable genes to calculate a diffusion map (29), which placed the distal cell population (cluster 15) at the root of a bifurcating cell fate trajectory into either phalanx (cluster 3) or interphalangeal joint progenitors (cluster 17) (Fig.S1B). Using Slingshot (30), we calculated pseudotemporal orderings of cells along these two diffusion map branches (Fig.S1C), resulting in a ‘phalanx’- and a ‘joint’-specific trajectory, respectively (Fig.1C). The top differentially expressed genes along these two trajectories contained many known markers of the respective cell populations, with their expected temporal expression dynamics (Fig.1D, Fig.S1D). For example, we were able to detect the expression of PFR markers, such as *INHBA* or *TCF7L2* (6, 31), at the onset of our pseudotime. Genes indicative of a progressive maturation of either phalangeal chondrocytes (e.g. *COL2A1, ACAN, MATN1*) or interphalangeal joints (e.g. *GDF5, BARX1, TGFB2*) were upregulated towards the respective ends of the two trajectories (32). Additionally, we identified novel markers and putative regulators of this bifurcating cell fate decision (Fig.S1D). Collectively, using scRNA-seq pseudotime analyses, we reconstructed the transcriptional dynamics at the PFR, documenting the expression dynamics of both known and novel marker genes, as digit progenitors differentiate into either phalanx or joint cell fates.

Given the genetic evidence supporting the importance of the TGF-beta superfamily in digit formation and patterning (26), we next focused on the transcriptional dynamics of all its members whose expression we detected in our HH29 sample. Hierarchical clustering based on expression dynamics across the trajectories revealed high temporal variance in six BMP genes, with three phalanx- and three joint-enriched signatures (Fig.1E). At the onset of our pseudotemporal progression, i.e. corresponding to the PFR and its immediate descendants, we detected *CHRDL1* and *BMPR1B*, whose expression extended into the phalanx branch, as well as *INHBA* (also known as *Activin Beta-A*), which additionally appeared re-activated later in the joint trajectory. After the bifurcation point of the phalanx and joint trajectories, *NOG* was transcribed in the phalanx trajectory and, to a lesser extent and with seemingly higher variability, in the joint trajectory. Also, *GDF5* and *CHRDL2* became expressed specifically in the joint progenitors, with *GDF5* being the only BMP ligand that we detected at appreciable levels in either of the two trajectories. These scRNA-seq expression profiles thus suggested that the majority of BMP activity during the phalanx-joint cell fate decision is driven by GDF5, signaling through its preferred receptor BMPR1B (24, 33).

Accordingly, we next investigated the spatiotemporal profiles of *GDF5* expression and BMP pathway activity *in vivo*. Using longitudinal tissue sections of chicken foot digit III at different developmental timepoints, we performed fluorescent *in situ* hybridization (FISH) for *GDF5* and combined it with fluorescent immunohistochemistry against the phosphorylated versions of SMADs 1, 5 and 9 (referred to as pSMAD thereafter), as a proxy for BMP pathway activity (Fig.1F-H). For both *GDF5* and pSMAD, we quantified normalized fluorescent intensities (NFIs) along the proximal-distal axis of the digit, and visualized them using either line plots or heatmaps (Fig.1I-N). This revealed repetitive peaks of restricted *GDF5* expression, corresponding to the forming interphalangeal joints, with broad shoulders of pSMAD activity marking the intervening phalangeal segments. The most distal digit domains, i.e. where full segmentation had yet to occur (‘Last joint to AER’), showed particularly dynamic profiles of *GDF5* and pSMAD. Counterintuitively, given its role as a BMP-activating ligand, wherever *GDF5* expression initiated, we observed a drop in pSMAD intensity, rendering the two activity profiles essentially out of phase from one another (Fig.1I, L). This occurs even though the entire distal digit domain expresses BMPR1B (6), i.e. all its cells should be competent to activate BMP signaling *via* GDF5. We therefore explored whether additional, potentially self-organizing mechanisms could explain the absence of pSMAD at sites of distal *GDF5* transcription, and the overall repetitive nature of observed BMP activity.

### Theory predicts that spatially periodic expression of the inhibitor *NOG* is required for digit patterning

Ectopic pSMAD activity is known to downregulate *GDF5* transcription in a cell-autonomous manner (34). Furthermore, GDF5 protein can diffuse and activate pSMAD at a distance (35). Therefore, GDF5 produced at the joint interzone could diffuse away to activate BMP signaling within the forming phalanx regions, where pSMAD would in turn downregulate further *GDF5* expression and thereby define antiphasic *GDF5/*pSMAD patterns. Collectively, these interactions delineate a long-range negative feedback loop, wherein diffusible GDF5 inhibits its own expression *via* activation of pSMAD at a distance. Such long-range inhibition is a hallmark feature of reaction-diffusion-based Turing patterns, with GDF5 fulfilling the requirements for a putative Turing inhibitor. In addition to GDF5 – an activator of BMP signaling – we reasoned that the reaction-diffusion system must incorporate an inhibitor of BMP signaling, to restrict pSMAD activity within the initiating joint interzone. Provided the inhibitor is secreted, this would also add a second diffusible factor to the proposed network, in addition to GDF5. Importantly, a model based on GDF5 and pSMAD alone – i.e. with only one diffusible species – could not form Turing patterns (15, 36). Indeed, of the six TGF-beta superfamily members that were dynamically expressed in our scRNA-seq pseudotime data, three were extracellular BMP inhibitors: *CHRDL1, CHRDL2*, and *NOG* (Fig.1E, (37)). Although overexpression of *CHRDL1* can cause loss of entire skeletal elements in the chicken hindlimb (38), no reported mutations in either of the two *CHRDL* genes affect joint patterning in mouse or human. In contrast, mutations in *NOG* are frequently associated with interphalangeal joint defects in humans, and *Nog(-/-)* mice fail to develop joints (26, 39). Therefore, we hypothesized that the diffusible BMP inhibitor NOG, together with GDF5 and pSMAD, would define a minimal Turing network capable of self-organized periodic patterning, to dictate the repetitive initiation of joint interzones.

To explore this hypothesis, we formulated a mathematical model of BMP signaling dynamics in the developing digit. Our core assumptions were that: (1) GDF5 activates pSMAD; (2) pSMAD inhibits *GDF5* transcription; (3) NOG binds GDF5 to form a complex; (4) the NOG-GDF5 complex cannot bind BMP receptors i.e., cannot activate signaling; and (5) NOG, GDF5 and the NOG-GDF5 complex diffuse in the extracellular space (Fig.2A). Initially, we also assumed *NOG* to be expressed uniformly throughout the digit ray, based on published expression data for mouse and chicken (11, 37, 39). Together these assumptions define a reaction-diffusion system, and we used partial differential equations (PDEs) to describe the spatiotemporal dynamics of each component (Fig.S2A, *SI Text S1*). Using linear instability analysis (36, 40, 41), we could derive general conditions that are necessary for periodic patterns to form as expected. Surprisingly, these analyses predicted that this reaction-diffusion system was unable to self-organize patterns for any combination of parameters. We therefore revisited the core assumptions of our model, focusing on the spatiotemporal dynamics of *NOG*, for which quantitative expression data had not previously been collected. We generalized the model and now allowed *NOG* expression to vary spatiotemporally, hypothesizing that feedback regulation of *NOG* expression by pSMAD activity may be required for the network to self-organize. When we re-analyzed our model with these more general assumptions, we were now able to identify a network that could spontaneously form patterns. We find that, regardless of model parameters, a necessary condition for patterning is that pSMAD activity must downregulate *NOG* expression (*SI Text S2*). Our analysis thus made the key prediction that *NOG* must be expressed in a periodic pattern, out-of-phase with pSMAD activity, for Turing-like pattern formation to occur (Fig.2B), contrary to previous reports of largely homogenous *NOG* expression along the digit (11, 37, 39). To investigate whether this prediction held true, we decided to compare the predicted spatiotemporal *in silico* dynamics of our Turing model with quantitative *in vivo* expression data.

**Figure 2.**
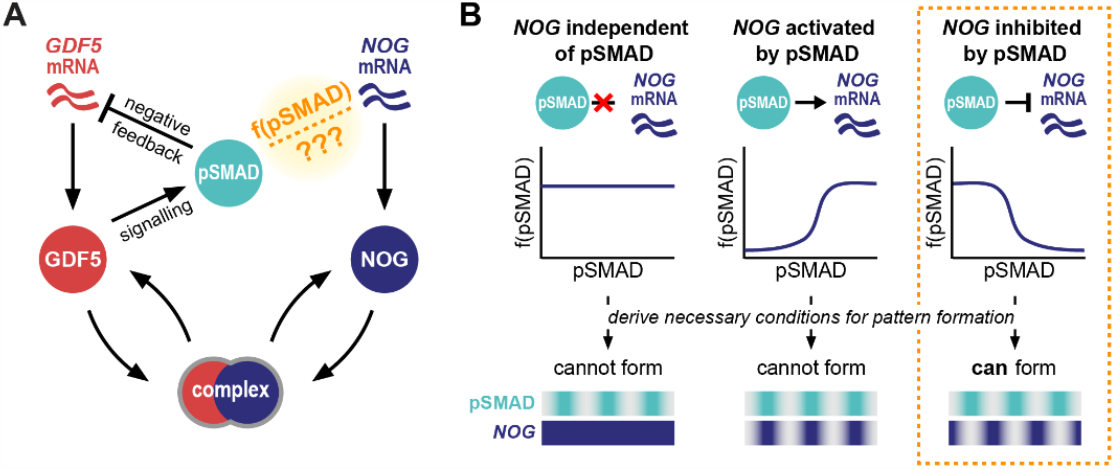
Theory predicts that BMP signaling must inhibit *NOG* expression for Turing-like joint patterning to occur. (**A**) Interactions in the BMP signaling pathway known to be involved in joint patterning: GDF5 activates BMP signaling (pSMAD) (69, 70); pSMAD negatively regulates *GDF5* transcription (34); NOG binds GDF5 to prevent it from signaling (71). It is not known how or whether active BMP signaling affects *NOG* transcription. (**B**) Mathematical modeling of this BMP network revealed that, regardless of biophysical parameters, self-organized periodic joint patterns can only form if *NOG* transcription is negatively regulated by active BMP signaling (see *SI Text S2* for proofs and linear instability analysis).

### A BMP-based Turing model recapitulates *in vivo* expression dynamics during digit patterning

We first simulated our model on a one-dimensional domain to understand its predicted dynamics, initially neglecting digit growth to focus on the intrinsic ability of the BMP network to self-organize. To satisfy the necessary condition for patterning derived above, we assumed that *NOG* expression was inhibited by BMP signaling (Fig.3A). We found that this network topology spontaneously formed periodic patterns with phase differences between the model components. To visualize these patterns, we plotted the concentrations of extracellular GDF5 and NOG protein, the level of pSMAD activity, and the inferred mRNA levels of *GDF5* and *NOG* (*SI Text S3*). We found that *GDF5* and *NOG* mRNAs were expressed in repeating peaks that were in-phase with one other, but out-of-phase with pSMAD activity. At the presumptive joint regions, where both *GDF5* and *NOG* are expressed, we predicted that high levels of NOG sequester GDF5 protein, thereby preventing it from activating the BMP pathway. However, at a farther distance from the joints, extracellular diffusion would allow unbound GDF5 to accumulate and activate BMP signaling, thus rendering the peaks of free GDF5 protein in phase with pSMAD (Fig.3B). We found that these self-organizing patterns form across a wide range of parameter space, with the same characteristic phase differences (Fig.S2B).

**Figure 3.**
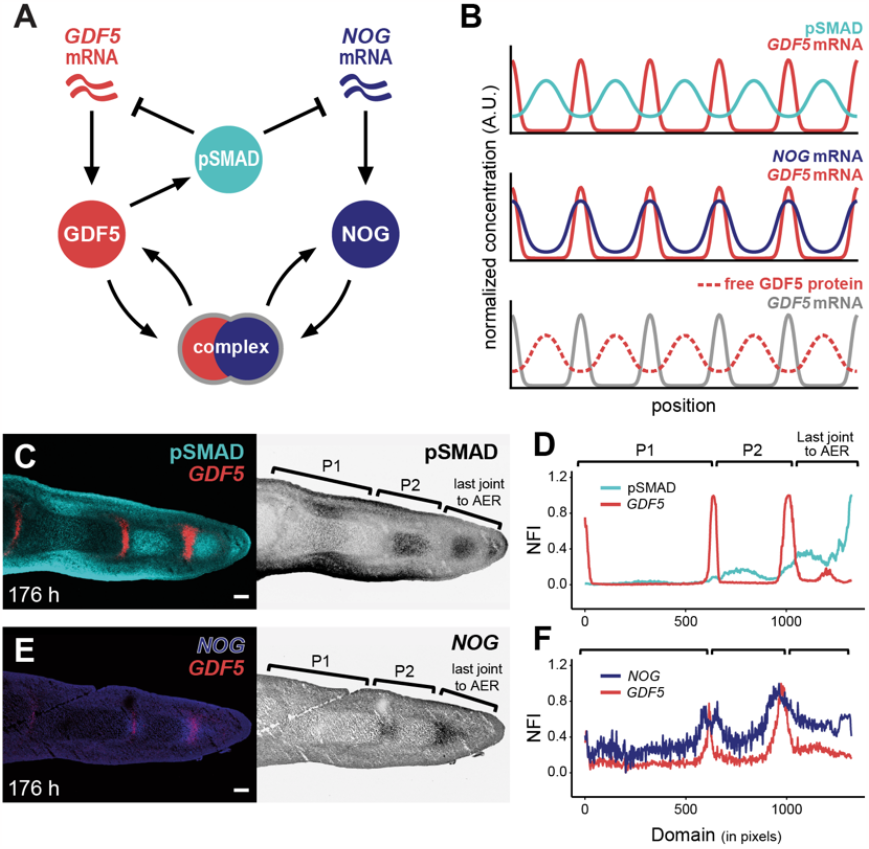
A BMP-based Turing model recapitulates *in vivo* signaling patterns and digit segmentation. (**A**) A BMP-based Turing model of digit patterning. (**B**) Simulated 1D expression patterns; plots show normalized expression of *GDF5* and *NOG* mRNA, pSMAD activity, and GDF5 protein dynamics along the proximal-distal digit axis. (**C**) Fluorescent RNA *in situ* hybridization for *GDF5* with immunohistochemistry for pSMAD on longitudinal sections of hindlimb digit III at 176h of development. (**D**) Plot of normalized fluorescence intensities (NFI) for pSMAD (cyan) and *GDF5* (red) (**E**) Multiple fluorescent RNA *in situ* hybridization for *GDF5* and *NOG* on an immediately adjacent section of **C**. (**F**) Plot of NFI for *GDF5* (red) and *NOG* (blue) along the proximal-distal domain of digit III at 176 h of development. Scale bars = 100 μm. 1 pixel = 1.243 μm.

To compare our *in silico* predictions to the spatiotemporal dynamics *in vivo*, we quantified expression patterns of *GDF5, NOG*, and pSMAD activity in the segmenting digits. We found that these *in vivo* measurements closely matched key predictions from our BMP-based Turing model. Namely, *GDF5* mRNA expression was observed in the presumptive joint regions, with periodic maxima of pSMAD concentration localized in the intervening phalangeal regions (Fig.3B-D). Double FISH, for *GDF5* and *NOG*, indicated that *NOG* mRNA expression is more dynamic than previously reported, forming an early repeating pattern of peaks in phase with *GDF5* mRNA (Fig.3E,F). Furthermore, by comparing the resulting profiles to quantifications of *GDF5* and pSMAD on adjacent sections, we confirmed that these distal *NOG* peaks were out-of-phase with pSMAD activity. In the proximal digit, corresponding to more mature digit segments, *NOG* expression continued to be expressed out-of-phase with pSMAD, but was excluded from the joint itself, and instead peaked at the borders of the forming interphalangeal joints (see Fig.3F [P1/P2 transition], Fig.S4C and (34)).

Whilst our model correctly predicted the relative phases of *GDF5, NOG* and pSMAD, so far it simulated patterning on a static domain. To explain the temporal progression of patterning observed within an elongating digit *in vivo*, we incorporated growth dynamics into our simulations, assuming that: (1) digits grow primarily at their distal tip, but also uniformly along their length; (2) cells irreversibly commit to either the joint or phalanx fate at a certain distance away from the PFR; and (3) patterning within the uncommitted distal digit domain is described by our BMP-based Turing model. These assumptions likely oversimplify more complex growth and differentiation dynamics; nonetheless, they provide insight into the potential impact of digit growth on phalanx-joint patterning. When we included growth and cell fate commitment in our simulations, we observed the same phase differences as before, but now, instead of forming simultaneously, *GDF5* peaks appeared sequentially. New joints formed towards the tip of the elongating digit, matching their iterative dynamics observed *in vivo* (Fig.S3A). Similar dynamics were observed for pSMAD activity, but with opposite phase, accompanying the periodic formation of sequential phalanges both *in silico* and *in vivo* (11) (Fig.S3A,B).

Importantly, the earliest evidence of symmetry breaking in the distal patterning domain closely matched our *in silico* predictions, with a drop in pSMAD activity coinciding with the initiation of *GDF5* mRNA expression (Fig.S3C,D, arrows). To quantify these patterns further, we combined measurements from multiple embryos and timepoints, initially restricting our attention to the distal end of the digit, to focus on the earliest patterning events that occur as cells exit the PFR. By aligning two consecutive peaks of *GDF5* expression in this distal domain, i.e., between the previously formed joint and the newly emerging one, we observed that multiple, superimposed profiles of *GDF5* expression and pSMAD activity were in near-perfect antiphase to one another (Fig.S4A). Moreover, *NOG* expression profiles all peaked around the newly forming joint, i.e. in phase with *GDF5* (Fig.S4A). This corroborated that the early, distal dynamics of patterning *in vivo* are consistent with the predictions from our BMP-based Turing model.

Quantifying expression profiles in more proximal digit regions, in which both phalanges-flanking joint interzones had already been specified, revealed alterations to the BMP signaling dynamics. Namely, profiles of both pSMAD and *NOG* evolved further, relative to the two flanking *GDF5* peaks marking the proximal and distal joints (Fig.S4B): the peak of the pSMAD shoulder shifted proximally, and its minimum no longer coincided with the distal joint, leading to a relative phase-shift and thus asymmetry between the *GDF5* and pSMAD patterns. *GDF5* expression at the distal joint increased, and its domain sharpened (Fig.S4B). Furthermore, the peak of *NOG* expression split, with the two resulting maxima now flanking the peaks of *GDF5*, as evidenced in embryos with slight developmental heterochronies between them (Fig.S4C). Importantly, however, these dynamics all occurred after the initial, segmentation-relevant symmetry breaking, and thus were not accounted for by our model.

Collectively, by combining *in vivo* data with *in silico* simulations, we found evidence for a BMP-based Turing system that underlies iterative phalanx-joint pattering during digit elongation. We also confirmed a key prediction of our model, namely that *NOG* should be expressed in a periodic pattern, out of phase with pSMAD.

### Spatiotemporal BMP signaling dynamics are evolutionarily conserved and predictive of known patterning perturbations

A repeating pattern of joints is a hallmark characteristic of digits across the tetrapod clade. Therefore, we wondered whether the self-organizing BMP network we identified in chicken could also be operating in other, distantly related species, to drive periodic joint patterns. To explore this, we examined BMP signaling dynamics during mouse digit development. We performed FISH for *Gdf5* combined with IHC for pSMAD on longitudinal digit sections at embryonic day E13.5, as well as double FISH for *Gdf5* and *Nog* on adjacent sections (Fig.4A,B). Line plot quantifications of normalized fluorescent intensities revealed that, indeed, similar expression dynamics were occurring during early mouse digit segmentation. Namely, while peaks of *Gdf5* expression were out of phase with pSMAD activity, *Nog* and *Gdf5* mRNA profiles were largely in phase with one another (Fig.4C,D). As in chicken, we observed a bimodal peak of *Nog* expression, centered on an already maturing *Gdf5*-positive joint interzone (Fig.4D), with the maximal levels of *Nog* being expressed out-of-phase with pSMAD (Fig.4, compare *Nog* to pSMAD in D and C). Peaks of pSMAD appeared to shift towards the proximal *Gdf5* bands (Fig.4C), although, based on our more detailed time series in the chicken, we hypothesize that this asymmetry is associated with phalanx maturation (see Fig.S4A,B). We thus concluded that the spatiotemporal dynamics of BMP pathway members during early digit patterning are largely conserved between birds and mammals. This supports the idea that the same core Turing system – involving GDF5, NOG and pSMAD – could be responsible for interphalangeal joint patterning across amniotes.

**Figure 4.**
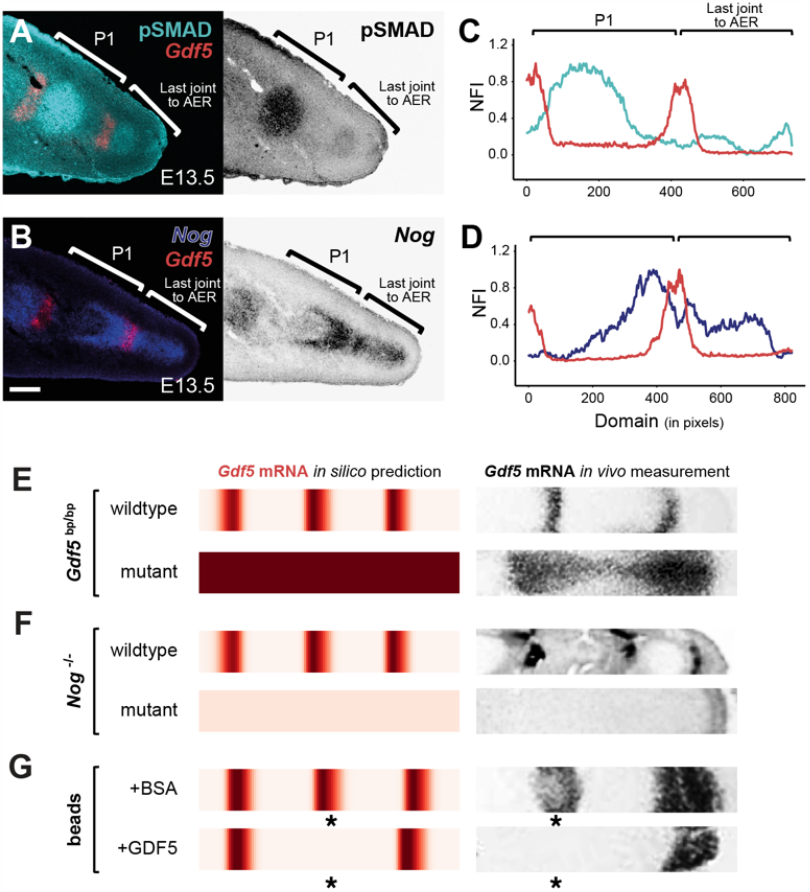
Conserved self-organized BMP signaling dynamics underlie mouse digit patterning and perturbation phenotypes. (**A, B**) Fluorescent RNA *in situ* hybridization for *Gdf5* combined with immunohistochemistry for pSMAD (**A**) and multiple fluorescent RNA *in situ* hybridization for *Gdf5* and *Nog* (**B**) on immediately adjacent sections of mouse digit 2 at E13.5. (**C, D**) Plots of normalized fluorescence intensities (NFI) of pSMAD (cyan) and *Gdf5* (red) (**C**) and *Gdf5* (red) and *Nog* (blue) (**D**) along the proximal-distal digit axis at E13.5. Scale bar = 100 μm. 1 pixel = 1.243 μm. **(E-G)** Heat map visualizations of *in silico* simulations to model *Gdf5* expression patterns in wildtype and *Gdf5*^*bp*^ (*brachypodism*) mutant backgrounds (**E**), in wildtype and *Nog* mutant backgrounds (**F**), and a wildtype background with beads soaked in either BSA or recombinant GDF5 protein, implanted at positions marked by an asterisk (**G**). Corresponding *in vivo Gdf5 in situ* hybridization measurements are provided at the side in black-and-white (images adapted from Storm and Kingsley, 1999^1^ for **E** and **G**, and from Brunet et al., 1998^2^ for **F**).

We then took advantage of molecular genetics studies in mice that have described joint patterning defects caused by mutations in the main components of our model. By changing model parameters, we tested whether mimicking genetic perturbations *in silico* would correctly predict the resulting effects on *Gdf5* expression, as an early marker of joint specification. First, we considered mutants in which *Gdf5* cannot activate BMP signaling. In this case, our model predicted *Gdf5* mRNA to be uniformly expressed at a high level throughout the simulated digit, instead of the characteristic repeating stripes. Such a pattern of *Gdf5* expression had indeed been reported in embryos carrying mutations disrupting the GDF5 coding sequence (Fig.4E, *brachypodism*) (23, 42), or lacking its essential signaling partner BMPR1B (43). Although this change in *Gdf5* transcription may appear counterintuitive – since inactivating an interzone marker led to an apparent expansion of the interzone domain – it is a natural consequence of the Turing-like negative feedback logic central to our model. Second, simulating mutants in which NOG is unable to inhibit GDF5 signaling activity predicted a downregulation of *Gdf5* expression and loss of its periodic pattern. Indeed, *Nog(-/-)* mutants fail to express *Gdf5* in digits, and do not develop interphalangeal joints (39) (Fig.4F). Finally, we considered a spatially restricted perturbation to digit patterning, by mimicking the effect of an implanted GDF5-soaked bead. This led to the local inhibition of *Gdf5* mRNA expression both *in silico* and *in vivo* (23) (Fig.4G, asterisks). Taken together, the ability of a single model to explain the expression dynamics and mutant phenotypes in both mouse and chick hints at a conserved BMP-based Turing mechanism that is responsible for digit patterning in birds and mammals, and perhaps more widely among the tetrapods.

### Segmentation and growth dynamics in two morphologically distinct digits

Phalangeal digit patterns display a large range of morphological diversity across the tetrapods, with the size and number of skeletal elements varying considerably both within and between species. We used our BMP-based Turing model to investigate the potential causes of these variations. As an *in vivo* model system to compare against, we took advantage of the chicken foot, in which each digit shows a distinct phalangeal formula. For example, in its adult form, digit III is longer than digit IV, but contains one fewer phalanx (Fig.5A).

**Figure 5.**
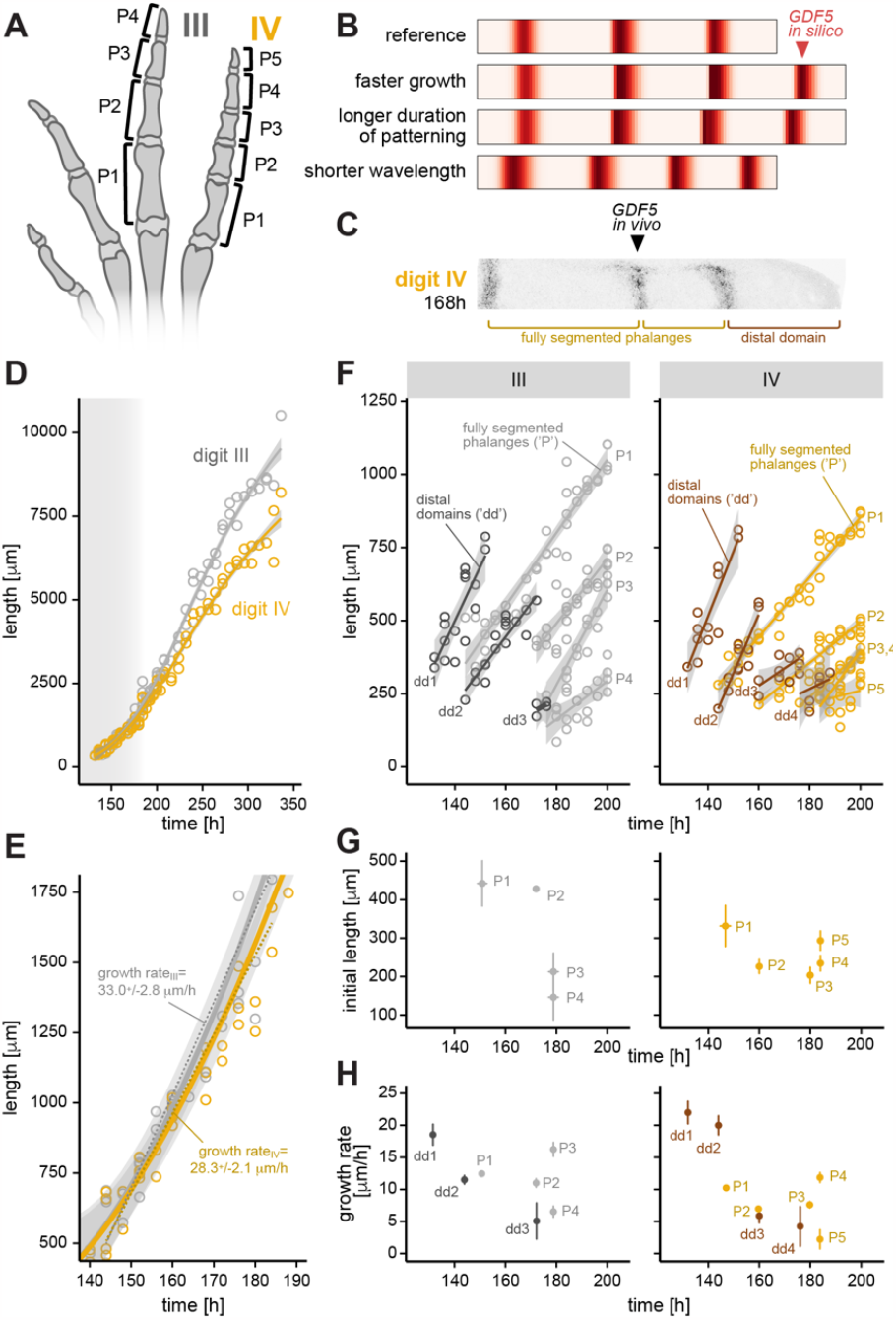
Growth and segmentation of two morphologically distinct digits of the developing chicken hindlimb. (**A**) Phalangeal morphologies of digits III and IV in the chicken foot. (**B**) *In silico* modeling predicts that an additional phalanx-joint element (= extra *GDF5* band) can form due to: i) faster growth, ii) longer duration of patterning phase, or iii) a shorter Turing wavelength. (**C**) Longitudinal section of digit IV at 168h, stained for *GDF5*. (**D**) Total digit growth dynamics for digits III and IV. Shaded grey area approximates the time window of digit segmentation. (**E**) Zoom-in of growth dynamics for digits III and IV during the phase of digit segmentation. A linear model of digit elongation for digits III and IV, from 144h to 184h, is superimposed to approximate the respective growth rates. (**F**) Growth and segmentation dynamics of individual elements for digits III and IV. Linear models of each element’s growth are plotted in either dark (dark grey, brown), for the distal domains (‘dd’) spanning from the last formed joint to the digit tip, or light colors (grey, yellow), for fully segmented phalanges (‘P’). (**G**) Initial mean length and developmental time of first appearance for the different phalanges of digits III (P1_III_-P4_III_; grey) and IV (P1_IV_-P5_IV_; yellow). (**H**) Initial growth rates of the different digit segments, either spanning from the last formed joint to the digit tip (dark: dark grey, brown), or fully segmented phalanges (light: grey, yellow) in digits III and IV.

We began by using our *in silico* model to explore different scenarios that could account for the observed differences in phalanx numbers. First, we found that increasing the total length of the segmenting digit domain, either through faster growth or a prolonged window for patterning, led to additional *GDF5* bands and hence the appearance of a supernumerary phalanx (Fig.5B, ‘faster growth’, ‘longer duration’). Alternatively, if the underlying wavelength of the Turing system was reduced, an additional skeletal element could be produced without changing digit length (Fig.5B, ‘shorter wavelength’). Indeed, we found that variations to several parameters in our model could result in significant changes to the pattern wavelength (Fig.S5A,B). For example, doubling the rate at which extracellular NOG was removed from the system sufficed to generate an extra phalanx *in silico* (Fig.5B, ‘shorter wavelength’).

To discriminate between these different scenarios, we produced quantitative growth and segmentation data for digits III and IV along a developmental time series (from 128h-336h; roughly HH27-HH40). Using longitudinal digit cryosections to identify consecutive bands of *GDF5* expression, we measured: the entire length of the digit, defined from the metatarsophalangeal joint to the AER; the lengths of individual, fully segmented phalanges; and the lengths of the unsegmented distal domain, defined from the most distal joint interzone to the AER (Fig.5C, see Materials and Methods for details). Looking at overall digit growth over time (Fig.5D), we saw that the major differences in digit lengths between digit III and IV only appeared after digit segmentation had completed (≈180h). During active digit segmentation (≈140-180h), growth rates of digits III and IV were nearly indistinguishable (Fig.5E). We thus excluded overall ‘faster growth’ as a possible explanation for the formation of an additional, fifth phalanx in digit IV.

To investigate potential digit-specific differences in pattering duration and/or wavelength, we next plotted the dynamics of each element of the two digits individually. Again, we discriminated between fully segmented phalanges (light colors) and the unsegmented distal domains (dark colors, Fig.5F). As expected, joint interzones initiated sequentially as the digit elongated to drive periodic phalanx segmentation (4). However, both within as well as across the two digits, the size, growth rates, and time of initiation varied between the different elements. Digit segmentation in digit III appeared to complete slightly earlier (before 180h) than in digit IV (after 180h) (Fig.5F,G). This difference in timing, however, appeared insufficient to fully account for the growth of an entire extra phalanx, compared to the durations seen for previously formed ones (≈15-20h, (Fig.5G)). We therefore investigated at which length each newly forming phalanx was individualized, i.e. was demarcated by the appearance of a faint *GDF5* band delineating its distal end (Fig.5G). This initial length serves as a proxy for the patterning wavelength in our *in silico* model, since it corresponds to the early patterning prior to subsequent growth. In both digits, distal phalanges showed a tendency to be progressively smaller at their onset, and were initiated at shorter time intervals. Importantly, however, up to the penultimate elements, phalanges in digit III (P1_III_, P2_III_) were longer at their point of initiation compared to their digit IV counterparts (P1_IV_, P2_IV_, P3_IV_). This suggested that the extra phalanx in digit IV may be, in part, attributed to a shorter wavelength of the underlying Turing system.

Finally, to explore the effect of growth on phalangeal proportions, we quantified the growth rates for each individual phalanx, as well as the unsegmented distal domains, and contrasted them across the two digits (Fig.5H). In both digits, most individual growth rates appeared to decline over developmental time, with the distal domains showing a steeper decline than the fully segmented phalanges. A notable exception to this trend were the two penultimate elements. In both digits, these phalanges (P3_III_, P4_IV_) showed a marked increase in their growth rates relative to the other digit elements. This highlights how variations to post-interzone initiation growth may influence the final morphology of the digit independently of early patterning.

Overall, our results reveal differences in growth and segmentation dynamics between two morphologically distinct digits. We find variations in phalanx size, both within as well as across the two digits, and show how these may be caused by alterations to phalanx-specific growth rates and/or to the reaction-diffusion parameters of the underlying Turing mechanism.

## Discussion

Digits within and across species vary in shape and size, due to changes in the numbers and the dimensions of their individual digit bones, the phalanges. Many of the morphological extremes relate to adaptations towards distinct modes of locomotion, such as the elongated phalanges in the digits of a bat wing, or the hyperphalangy with more than 15 individual skeletal elements per digit in the flippers of certain whales (4, 44). Here, we have studied the early developmental basis for this morphological diversity, focusing on the molecular and cellular dynamics giving rise to distinct digit segmentation patterns. Combining quantitative *in vivo* data with *in silico* simulations, we present a model in which the segmentation wavelength is determined by a Turing-like mechanism involving the BMP-ligand GDF5 and its extracellular inhibitor NOG. Furthermore, distinct growth rates for individual phalanges, both within and between digits, seem to have enabled a highly modular approach to diversifying digit morphologies over the course of tetrapod evolution.

Based on pseudotemporal ordering of scRNA-seq transcriptomic data and *in vivo* quantifications of BMP signaling dynamics, we propose GDF5 – a BMP-activating ligand expressed and secreted at forming joint sites – as the inhibitor of a Turing-like reaction-diffusion system. Furthermore, theory and *in vivo* data suggest that highly dynamic *NOG* expression, in phase with *GDF5*, completes this early self-organizing process to specify joint interzone locations. By modeling the reaction-diffusion dynamics of *GDF5* and *NOG in silico*, we predict that *NOG* must be expressed out-of-phase with pSMAD – the intracellular effector of active BMP signaling – for the system to organize into repetitive patterns. While this notion was at odds with previous *NOG* expression studies, it was nonetheless confirmed by our quantitative FISH measurements. Importantly, we find that the transcriptional signatures and BMP signaling dynamics predicted by our model closely match quantitative expression data from developing digits, in both chicken (Fig.3) and mouse (Fig.4). Moreover, the model successfully phenocopies aberrant digit patterns resulting from perturbations to the BMP pathway in mouse. Similar digit malformations are observed in humans with analogous genetic alterations, with *GDF5, NOG*, or *BMPR1B* commonly associated with digit patterning defects (26). Our model can therefore explain the etiology of congenital human conditions in which the digits lack either interphalangeal joints (symphalangism (45)), or phalanges (Chondrodysplasia Grebe type (46)). Collectively, these results thus suggest that a conserved BMP-based Turing mechanism is responsible for digit segmentation patterns across the amniotes.

An intriguing feature of our model is its simplicity, with the initial segmentation patterns driven by a single developmental signaling pathway. Undoubtedly, specifying distinct digit morphologies *in vivo* will involve multiple other signaling pathways, as well as more complex morphogenetic processes than we have approximated *in silico*. These are likely to include changes in cell shape and adhesion, integration of mechanical forces, and differential regulation of long bone growth (9, 47, 48). However, by considering the BMP pathway alone, we were still able to capture the earliest expression dynamics associated with digit segmentation and predict key mutant phenotypes. Moreover, genetic evidence from the mouse suggests that other signaling pathways known to affect digit patterning are likely dispensable for the periodic placement of the joint interzones (e.g., WNTs (49, 50), Hedgehog signaling (11, 51), extradigital BMPs (52, 53)), suggesting that interactions between GDF5, NOG, and pSMAD may be sufficient for the initial symmetry breaking. A further simplification of our model is that we consider patterning only within a 1D domain, rather than the full 3D geometry of the developing digits. The predictions in Figure 2 hold regardless of dimension, meaning our model will form periodic patterns with the same overall phases in 2D or 3D as we here presented for 1D. Nevertheless, beyond the earliest symmetry breaking events, we expect that other pathways will be involved in refining a 3D digit pattern. In particular, we hypothesize that while our BMP-based model can initiate repetitive patterns with periodic dots of pSMAD activity, other pathways – or BMP signaling components – will be required to sharpen the initially broad domains of *GDF5* into straight, narrow stripes (12). Hedgehog signaling is a promising candidate for future investigation, since *Gdf5* bands remain broad and curved in *Gli3*(-/-) digits (11). Furthermore, *CHRDL1* and *CHRDL2*, with their complementary patterns in the phalanx and joint domains, respectively, may also contribute to refine the stripe-like pattern of late *GDF5* expression (Fig.1E). And finally, whilst *BMPR1B* is expressed throughout the distal digit (6), thus rendering a ligand-receptor-based Turing mechanism unlikely (54), subsequent regulation of *BMPR1B* levels may contribute to the maturing *GDF5* pattern. Indeed, the exclusion of *BMPR1B* expression from the maturing joint itself, and the importance of *BMPR1A* at later stages, may reflect a transition into a new BMP signaling regime (Fig.1E, (6, 55, 56)).

*NOG* and pSMAD both show changes in their expression and activity patterns that are likely related to a switch in the predominant mode of phalanx elongation, transitioning from mostly distal progenitor proliferation to epiphyseal plate-driven long bone growth. In proximal phalanges, we see a phase shift of the pSMAD pattern relative to the *Gdf5* peaks, with the pSMAD peaks moving towards the proximal joint, and an increase of pSMAD being observed near the distal joint (Fig.S4B). During long bone development, two peaks of pSMAD activity mark the distal growth zones at either end of the skeletal element. These pSMAD domains rely on BMPs from the perichondrium and the epiphyseal plate itself and are essential for signaling crosstalk and endochondral bone elongation (57). Furthermore, two defined bands of NOG flanking the interzone safeguard continuing joint formation against these BMP signals (34, 58, 59). This split in the early interzone-centered domain of *NOG* expression seems to evolve as the forming phalanges transition from distal domain segmentation to long bone growth (Fig.S4B,C).

Like for the observed BMP signaling dynamics, the presence of two distinct temporal regimes – i.e. pre- and post-segmentation – also manifests itself in digit growth, both within and across digits. Changes in total digit lengths within the same autopod appear to largely arise during the post-segmentation phase (Fig.5D). Before that, however, growth rates already differ amongst individual phalangeal elements within a digit, highlighting the modular nature of digit elongation (Fig.5F-H, (16)). For most phalanges, these growth rates progressively decline as development proceeds (Fig.5H). This occurs to a similar extent in both digits III and IV, and likely relates to the waning of FGF signals from the AER, and – potentially – its impact on BMPR1B transcription in the distal patterning domain (10, 60). Strikingly, however, once the penultimate phalanges have formed, they show a pronounced up-tick in growth (P3_III_, P4_IV_; Fig.5H). While the molecular underpinnings remain unknown, nature seems to have exploited this modular mode of growth regulation in the evolution of elongated, penultimate phalanges in the feet of raptors (16). Specification of the last phalanx – known to develop differently from the rest (10, 61) – might additionally alter the segmentation dynamics at the distal end of the digit.

Comparing across digits, the elements in digit IV initiate at a shorter wavelength than in digit III, for all phalanges up to the penultimate one (Fig.5G). It is tempting to speculate that variations in BMP activity across the anteroposterior axis of the limb might provide a mechanistic link between our Turing model and morphologically distinct digit identities (7, 25, 62, 63). For example, BMPs from the interdigital mesenchyme may diffuse into the distal digit where they can then bind and thereby remove NOG from the system, a scenario which is predicted to decrease the segmentation wavelength (Fig.5B, Fig.S5B). However, regulatory interactions between mesenchymal BMPs and the AER, at the distal margin of the autopod, might make it difficult to experimentally discriminate between changes in segmentation wavelengths and altered digit elongation dynamics (62, 64). Ultimately, though, the most extreme variations in digit morphologies have arisen between different species, from either alterations in post-patterning growth – such as the elongated digits in bats (65) – or a prolonged patterning window, as proposed for cetacean hyperphalangy (64, 66). The observed differences in segmentation wavelengths in our chicken data might thus not be due to any particular adaptive trait. Rather, they may relate to an ancient developmental constraint, due to molecular interactions between the anterior-posterior patterning system of the limb and the BMP signaling pathway (11), which manifested itself already in the ancestral condition of tetrapod digit formulas (67, 68).

Collectively, our work has identified a conserved BMP-based Turing system that is involved in the formation of the repetitive joint segmentation pattern that characterizes tetrapod digits. Furthermore, by combining quantitative *in vivo* data with *in silico* simulations, we explore how variations in growth and patterning dynamics can give rise to highly distinct digit morphologies, and demonstrate how modulation of a self-organizing segmentation wavelength can alter the number of individual elements – here, the phalanges – in a repetitive pattern.

## Materials and Methods

### Pseudotime analyses

Pseudotime analysis was performed using a previously published autopod HH29 single-cell RNA-sequencing data set (see (27) for details on quality checks, normalization and cluster identification using *Seurat* v3.1.4 (72)). Three skeletogenic clusters (cls. 3, 15 and 17) were used for further analyses. Using highly variable genes, a diffusion map of the three clusters was calculated with the R package *Destiny* (29). Calculation of pseudotime trajectories was performed with *Slingshot* (30). Finally, differential expression analysis along the pseudotime was done using the *MAST* package (73), and expression heatmaps were visualized in *R* studio.

### Embryo tissue sampling and processing

Fertilized chicken eggs were incubated at 38°C in a humified incubator and harvested and staged according to the Hamburger-Hamilton developmental table (28). Mouse embryos were isolated and processed for analysis by M. Luxey in accordance with national laws and experimental procedures approved by the Regional Commission on Animal Experimentation and the Cantonal Veterinary Office of the city of Basel (license 1951 to Rolf Zeller and Aimee Zuniga). Embryos were dissected in ice-cold PBS and fixed with 4% paraformaldehyde for 2 h on ice. For cryosections, digit tissue was dehydrated in a sucrose gradient up to 30% sucrose/PBS and cryopreserved in OCT (Leica). Cryosectioning was performed at 18 μm thickness.

### Immunohistochemistry on cryosections

Cryosections were air dried for 15 min at room temperature (RT) and washed in Tris-buffered saline (TN: 0.1 M Tris pH 7.5, 0.15 M NaCl). For pSMAD, a step of antigen retrieval was added (1:5000 Proteinase K in TN for 10 min at RT), followed by post-fixation for 5 min in 4% PFA. Endogenous peroxidase was inactivated with 0.3% H_2_O_2_ in TN for 1h at RT. Slides were blocked in 0.5% BR (Blocking Reagent (Akoya Biociences))/TNT (TN, 0.05% Tween) for 1h at RT and incubated with a primary antibody against pSMAD1,5,9 (Cell Signaling 13820S, rabbit, 1:300) overnight at 4°C. Slides were washed 3 times in TNT and incubate with a biotinylated secondary antibody, followed by streptavidin-conjugated peroxidase incubation and signal amplification using the TSA Plus Cyanine-3 or -5 kits (Akoya Biosciences).

### Fluorescent *In situ* hybridization on cryosections

Probes were produced *in vitro* transcription using T3, T7 or SP6 RNA polymerases (Promega) and digoxigenin or fluorescein labeled nucleotides (Roche). *In situ* hybridization was carried out using standard protocols (74), incubating with either anti-digoxigenin or anti-fluorescein-POD antibodies (Roche) diluted at 1:300. Signal amplification was done with 1:50 TSA Plus Cyanine-3 or -5 (Akoya Biosciences) for 1 h at RT. For dual-probe RNA *in situ* hybridization, an inactivation step after the first TSA Plus incubation was performed in 0.6% H_2_O_2_/TNT for 1 h at RT. Fluorescent *in situ* hybridization combined with pSMAD staining was performed first, followed by immunohistochemistry in TNT buffer, as described above.

### Measurements of normalized fluorescence intensities in developing digits

Fluorescent signals were imaged with a confocal microscope (*Olympus Fluoview FV3000*). Pictures were stitched with the *Pairwise-stitching* plug-in in *Fiji*. After image processing, average fluorescence along the metacarpal to AER axis was measured in *Fiji*, by quantifying pixel intensities along a line width of 100 pixels (‘Analyze>Plot profile’). Fluorescence intensities were scaled from 0 to 1, based on maximum and minimum values. Finally, plots of normalized fluorescence intensities (NFI) were visualized as line plots or heatmaps in *R* studio.

### Mathematical modeling

Full details of the modeling are provided in the theory supplement. Briefly, we constructed a reaction-diffusion model of BMP signaling in the developing digit. We used systems of partial differential equations (PDEs) to describe these dynamics, with molecular interactions as schematized by Fig 2 or Fig 3A. These PDEs were first analyzed using linear instability analysis to derive necessary conditions for pattern formation to occur. We then performed 1D simulations using custom MATLAB scripts. For further details of simulation methods and model parameters, please see *SI Text S3*.

### Length measurements of individual digit elements

Chicken hindlimbs were collected along a developmental time series: every 4h, from 128h to 204h of development, and every 8h, from 204h to 336 h. Up to two embryos were collected per time point and both hindlimbs of each embryo were dissected and analyzed. Chromogenic or fluorescent *in situ* hybridization for *GDF5* was performed on early stages, and simple DAPI stains on later stages. Slides were imaged and pictures stitched with the *Pairwise-stitching* plug-in in *Fiji*. Length measurements of individual elements were performed in *Fiji*, by drawing a line from the middle of the proximal joint (as defined by *GDF5* signal, or a gap in the DAPI channel) to the middle of the distal joint, or from the most distal joint to the AER. For total digit length measurements, lengths of all individual elements were summed up. Plots of the digit and phalanx growths were done in *R* studio.

## Author Contributions

This study was conceived and designed by EG, TWH and PT. Pseudotime analysis was conducted by CF and PT. Experimental embryology, stainings, imaging and analyses were carried out by EG, with help from SF and ADC. Mathematical modeling was carried out by TWH. EG, TWH, and PT wrote the paper, with feedback from the other authors.

## Acknowledgments

The authors wish to thank C.J. Tabin, E. Clark and J.C. Scoones for a critical reading of the manuscript, M. Luxey, A. Zuniga and R. Zeller for providing wild-type mouse embryos, F. Sacher and M. Wang for help with *R*, D. Barac for conceptual input on the developmental digit growth series, and D. Ebert and all members of our groups for useful discussions. Calculations for scRNA-seq analyses were performed at sciCORE (http://scicore.unibas.ch/), scientific computing center at the University of Basel. This work was supported by research funds from the UKRI Biotechnology and Biological Sciences Research Council [grant numbers BB/W003619/1 and BB/X511973/1] to TWH, and from the Swiss National Science Foundation [SNSF project grant number 310030_189242] and the University of Basel to PT.

## Supplementary Figures

**Figure S1.**
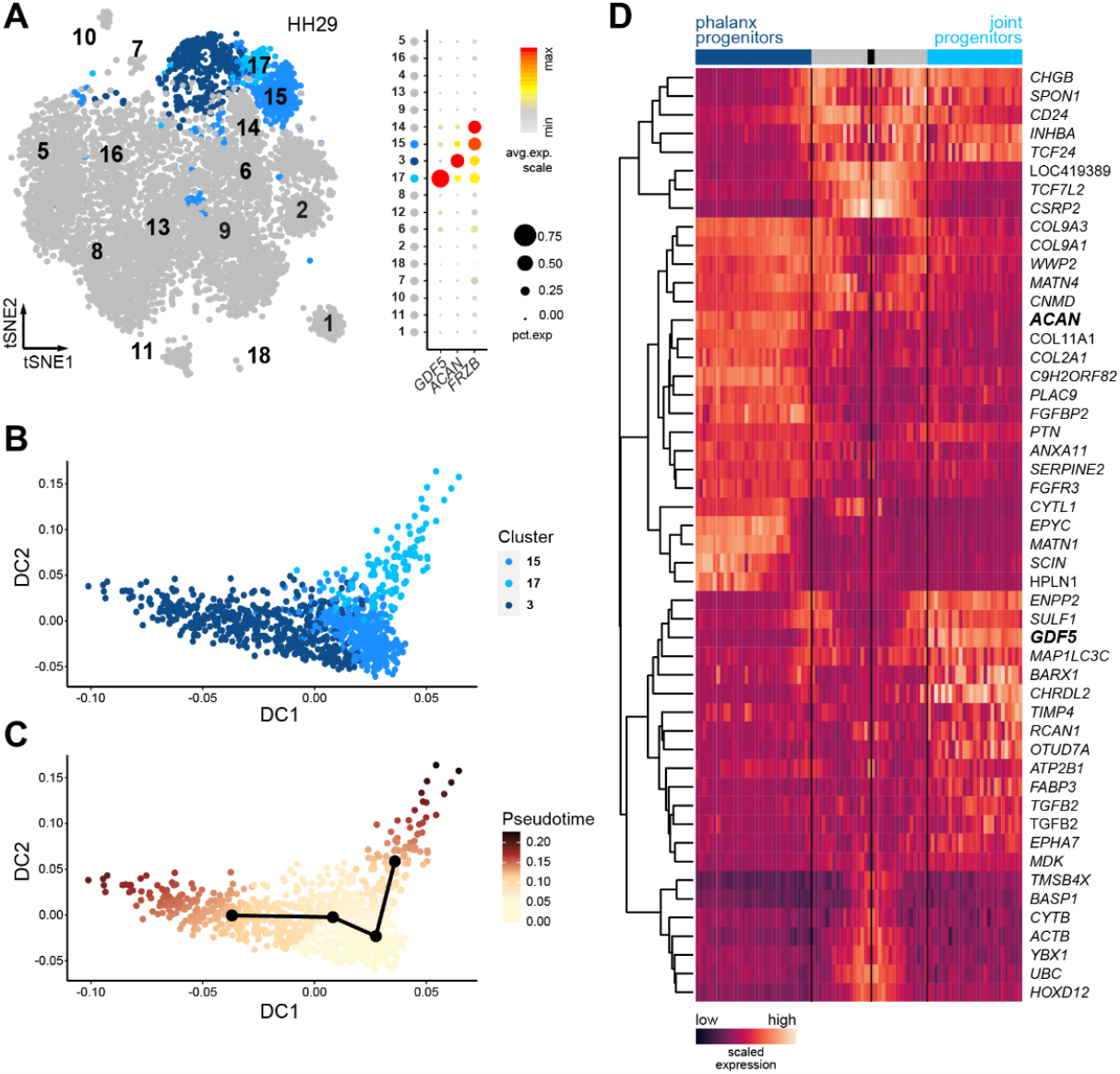
Single-cell pseudotemporal reconstruction of a bifurcating cell fate decision into phalanx or joint progenitor cells. (**A**) tSNE representation of a HH29 chicken hindlimb scRNA-seq dataset. Based on marker gene expression, cluster 3, 15 and 17 were selected for pseudotime analysis, to represent maturing phalanx progenitors (3), naïve skeletal progenitors and mesenchymal cells (15) and joint progenitors (17), respectively (see Feregrino et al., 2019 for details). (**B, C**) Diffusion map of the three clusters and their relative cellular contributions (**B**), as well as overall pseudotime progression along two bifurcating trajectories (**C**). (**D**) Pseudotime heatmap of top differentially expressed genes between the phalanx progenitor trajectory (dark blue) and the joint progenitor trajectory (turquoise). The black box corresponds to the starting point of the pseudotime, the grey zone to the part shared by the two trajectories. Scaled gene expression from low (purple) to high (orange). Non-italic names correspond to proteins identified by manual *blastx* using the cDNA sequences of the respective ENSGAL IDs.

**Figure S2.**
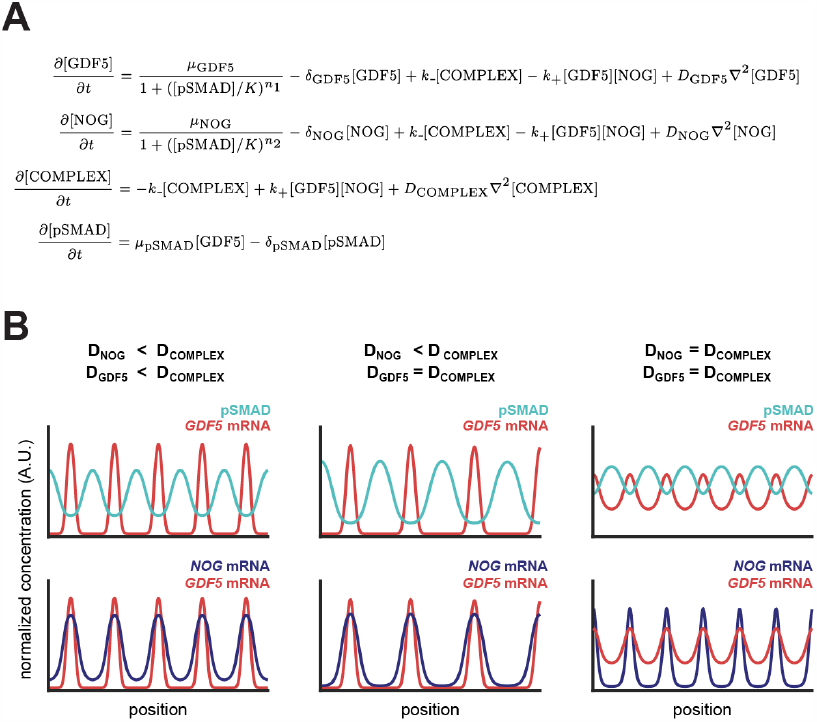
A mathematical model of self-organized BMP signaling in the developing digits. (**A**) Partial differential equations describing the spatiotemporal dynamics of BMP signaling in the developing digits. See *SI Text S1* for more details on the model. (**B**) Many different parameter combinations generate self-organized periodic patterns *in silico*. Here are shown three example parameter sets, with rapid complex diffusion (left); slow NOG diffusion (middle); and no differential diffusivity (right).

**Figure S3.**
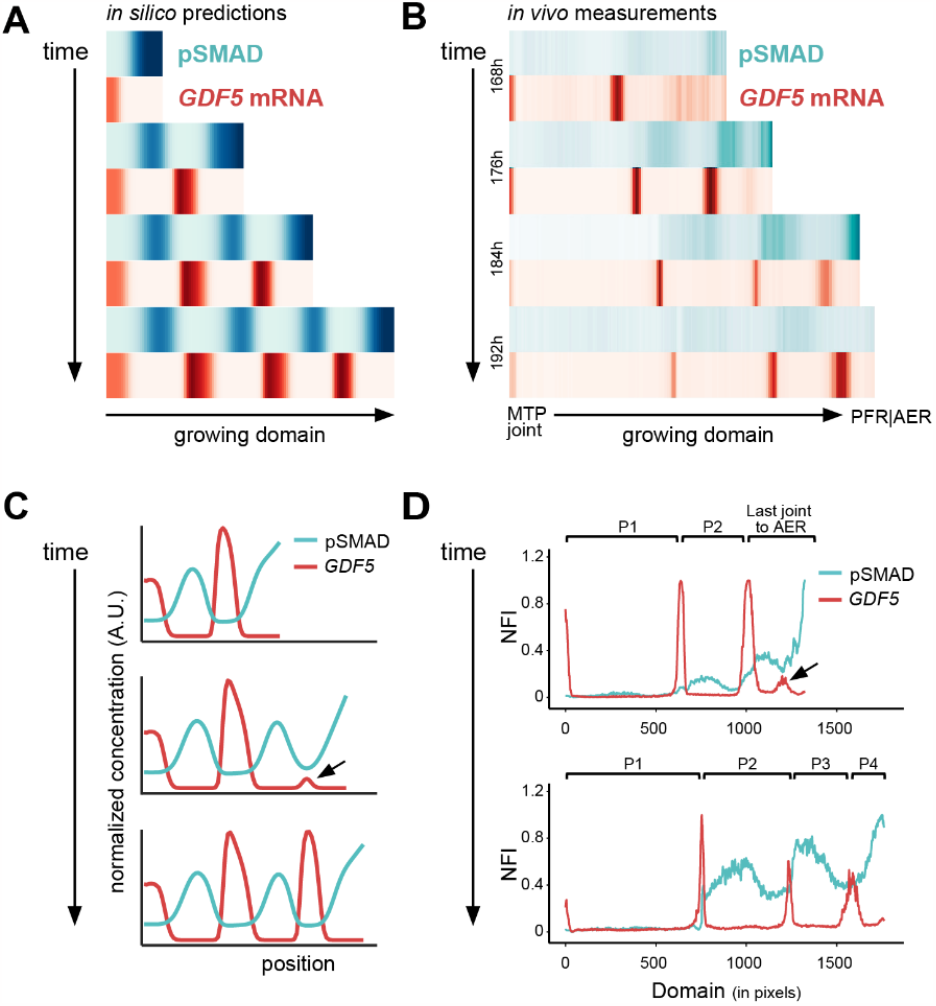
BMP signaling dynamics and patterns during digit development. (**A**) Heat map visualizations of model simulations of normalized pSMAD intensities (top, cyan) and normalized *GDF5* expression (bottom, red), along the proximal-distal axis of a simulated growing digit domain at different timepoints of development. (**B**) Heat map visualization of normalized *in vivo* fluorescence intensities of pSMAD immunohistochemistry (top, cyan) and *GDF5 in situ* hybridizations (bottom, red) along the proximal-distal axis of digit III at different indicated timepoints of development. The very distal zone of high pSMAD corresponds to the PFR. (**C, D**) Temporal progression of pSMAD and *GDF5* dynamics *in silico* (**C**) and *in vivo* (**D**). Weak *GDF5* expression initiates in the distal digit where pSMAD is downregulated, marking the future site of the forming joint (see arrows in **C, D**).

**Figure S4.**
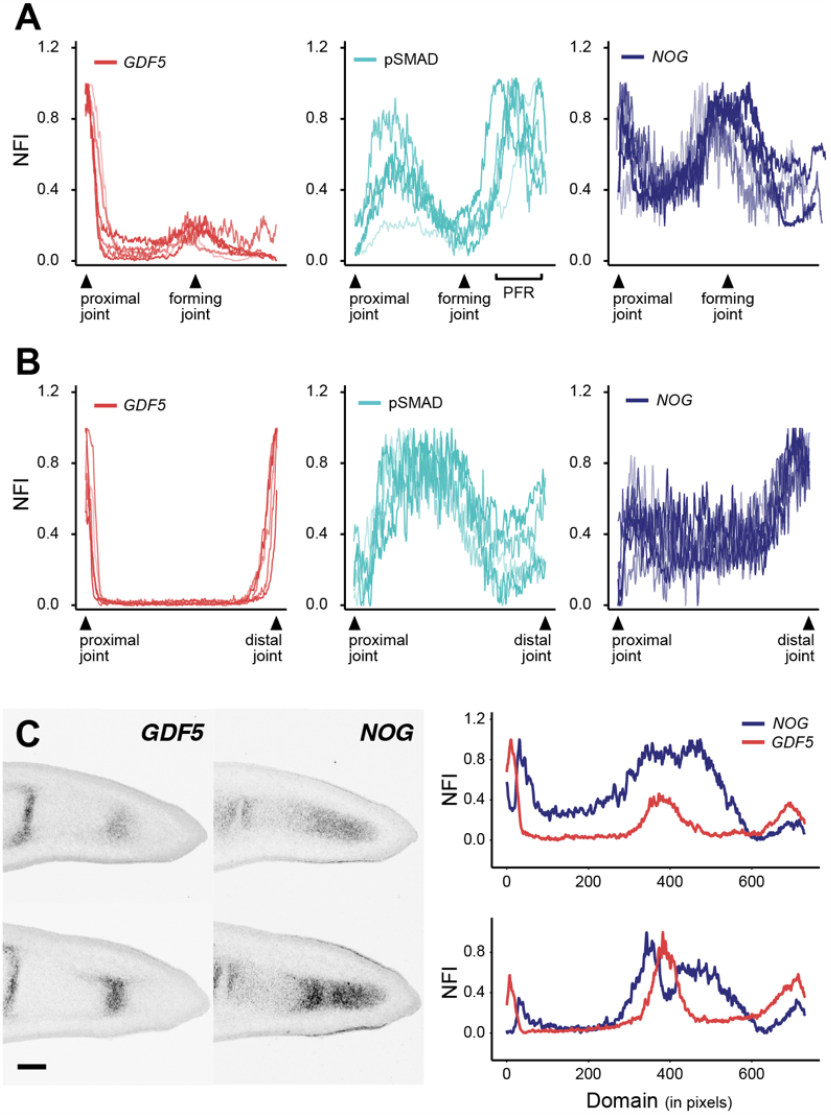
*In vivo* BMP signaling dynamics of forming and fully segmented phalanges. (**A**,**B**) Quantification of expression dynamics in newly forming (A) and already segmented (B) phalanges. (**A**) Superimposition of multiple normalized fluorescence intensity (NFI) curves for *GDF5* (red), pSMAD (cyan) and *NOG* (blue) in newly forming phalanges. The region from the most distal joint to the digit tip was measured, and individual lengths adjusted to align the corresponding peaks of *GDF5* expression. (**B**) Superimposition of multiple normalized fluorescence intensity (NFI) curves for *GDF5* (red), pSMAD (cyan) and *NOG* (blue) in fully segmented phalanges. The region spanning two consecutive *GDF5* peaks were measured, and individual lengths were adjusted to align the *GDF5* peaks. (**C**) Spatiotemporal variations in *GDF5*/*NOG* patterns in newly forming phalanges. The two images are from embryos collected at the same timepoint, with the lower showing signs of a slightly more developed distal domain. Distal patterns of *NOG* are in phase with *GDF5* expression, but as the newly formed phalanges mature and upregulate *GDF5*, the *NOG* domain becomes split into two *GDF5*-adjacent peaks.

**Figure S5.**
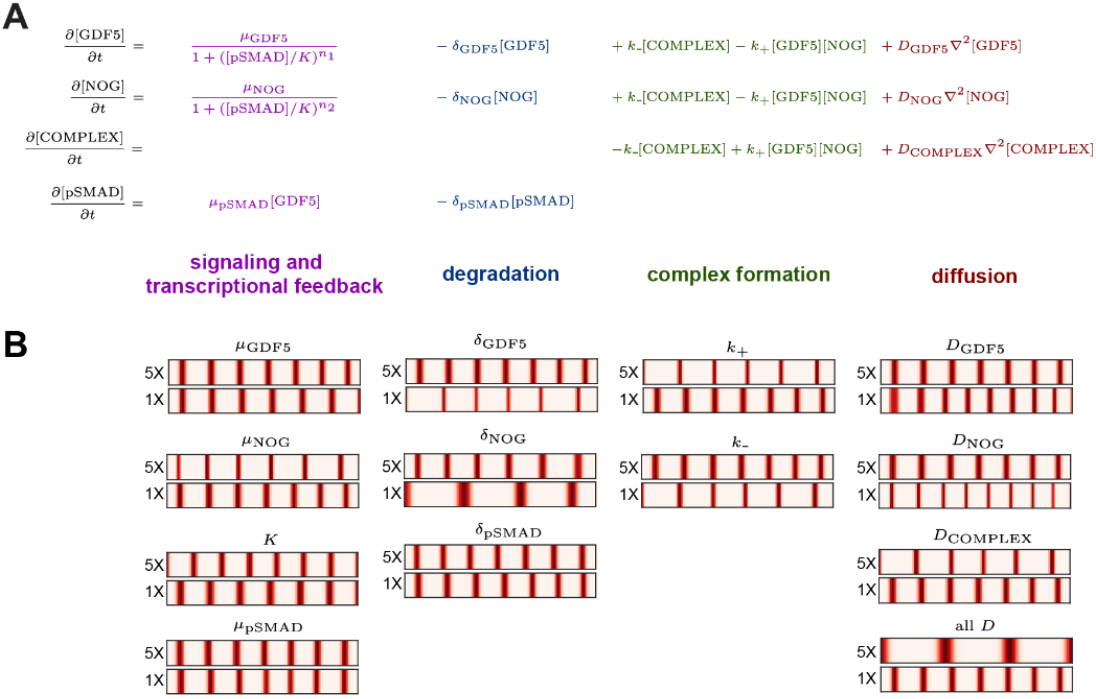
Examples of parameter variations that change pattern wavelength *in silico*. (**A**) Partial differential equations of the BMP-based Turing model, with terms and parameters colored according to the molecular processes they represent (**B**) Model parameters were individually varied across a five-fold range and simulations performed to predict the effect on pattern wavelength. We find that multiple parameters significantly impact pattern wavelength. For the final perturbation (all D, lower right), the diffusion constants for GDF5, NOG and COMPLEX were all varied by the same amount.

## Supplementary Information

### S1. Model formulation

We describe the spatiotemporal dynamics of BMP signaling in the digits by the following system of partial differential equations (PDEs):

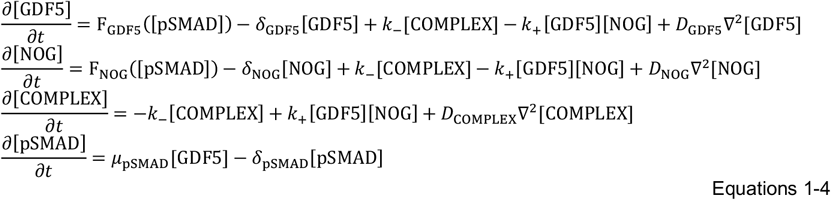

Here, […] denotes the concentration of each molecular species, which varies in space within the digit and in time as development proceeds. The processes which define the dynamics of this system are:

#### Transcription/secretion

We assume that GDF5 and NOG are produced and secreted by cells within the digit, at rates F_GDF5_ and F_NOG_ respectively. Given that active BMP signaling inhibits *GDF5* transcription, we choose F_GDF5_ to be a decreasing function of pSMAD. We also consider the possibility of feedback on *NOG* transcription by allowing F_NOG_ to vary with pSMAD.

#### Complex formation

We assume that extracellular GDF5 and NOG reversibly form a complex: GDF5 + NOG ↔ COMPLEX, with an on-rate *k*_+_ and an off-rate *k*_−_. We use mass action kinetics with a 1:1 stoichiometry (Groppe et al., 2002; Schwaerzer et al., 2012).

#### Signaling

We model BMP signaling via the dynamics of pSMAD, assuming that pathway activation increases linearly with the concentration of GDF5 ligand.

#### Degradation

We model the degradation of GDF5 and NOG in the extracellular space with rates *δ*_GDF5_ and *δ*_NOG_ respectively, and the intracellular turnover of pSMAD with rate *δ*_NOHP_.

#### Diffusion

We allow all extracellular species (GDF5, NOG, and complex) to diffuse with rates *D*_GDF5_, *D*_NOG_, *D*_COMPLEX_ respectively.

##### Notation

Here we adopt the convention that capitalized variables refer to proteins (e.g., GDF5), whereas italicized variables refer to mRNA gene expression (e.g., *GDF5*).

### S2. Deriving necessary conditions for pattern formation

Numerous experimental observations suggest that a Turing instability is responsible for the repetitive patterning of joints within the digit (Scoones & Hiscock, 2020). We therefore investigate the conditions that would allow Equations 1-4 to form Turing patterns.

A necessary condition for Turing instabilities is that diffusion causes the homogeneous steady state to become unstable with respect to spatially periodic disturbances. Using well-developed tools for linear instability analysis (Diego et al., 2018; J.D. Murray, 2008), we begin by computing the steady states of Equations 1-4 in the absence of diffusion. We denote the steady state levels of {GDF5, NOG, complex, pSMAD} as {*G*_0_, *N*_0_, *C*_0_, *S*_0_} respectively, and find that:

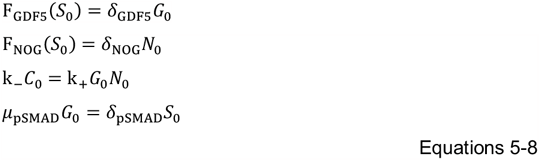

Next, we explore the response of this homogeneous steady state to small perturbations. We define new variables {Δ*g*, Δ*n*, Δ*c*, Δ*s*} that describe the normalized deviation of concentrations about the steady state for {GDF5, NOG, complex, pSMAD} respectively (e.g., Δ*g* ≡ ([GDF5] − G_0_)/G_0_). Substituting into Equations 1-4, and keeping only the first order (linear) terms, we obtain:

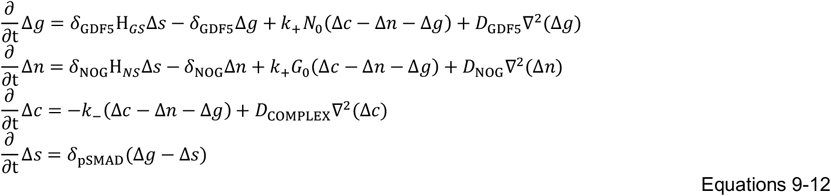

Here, the H_*ij*_ terms refer to the normalized sensitivities of the transcriptional functions. Specifically, the term:

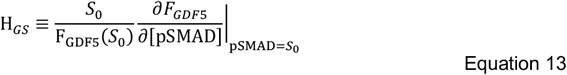

captures how the transcription of *GDF5* depends on the level of pSMAD; we know that *H*_*GS*_ < 0 since pSMAD inhibits *GDF5*. Similarly,

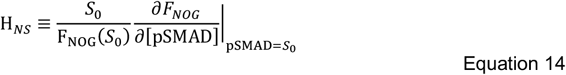

describes the potential feedback between pSMAD activity and *NOG* transcription. If, as published data suggests (Brunet et al., 1998; Huang et al., 2016; Lorda-Diez et al., 2013), *NOG* is expressed uniformly along the digit then *H*_*NS*_ = 0. If pSMAD activates *NOG*, then *H*_*NS*_ > 0; if pSMAD inhibits *NOG*, then *H*_*NS*_ < 0.

To examine the instability of the steady state to periodic disturbances, we apply the Fourier transform to Equations 9-12, yielding:

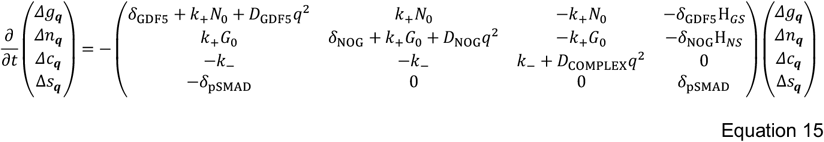

We refer to the reaction-diffusion matrix on the right hand side of Equation 15 as **F**^**RD**^(*q*^2^), mirroring the notation from (Diego et al., 2018).

A necessary condition for a diffusion-driven instability is that det[−***F***^***RD***^(*q*^2^)] must become negative for some positive value of *q*, whilst remaining positive at *q* = 0. Evaluating this determinant gives:

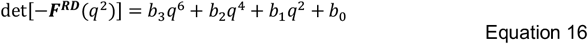

where the coefficients {*b*_*i*_} may be expressed in terms of the parameters in Equation 15. (The algebraic manipulations are computed using the Symbolic Math Toolbox in MATLAB). Inspecting these coefficients, and recalling that *H*_*GS*_ < 0, we immediately see that *b*_3_ > 0, *b*_2_ > 0 and *b*_0_ > 0 for all possible model parameters. Using Descartes’ rule of signs, we thus find that *b*_1_ < 0 is a necessary condition for the polynomial in Equation 16 to change sign for some positive value of *q*. Explicitly writing the parametric dependence of *b*_1_ gives:

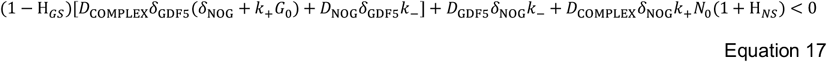

Since we know that *H*_***GS***_ < 0, this inequality can only be satisfied if the final term is negative, i.e.,

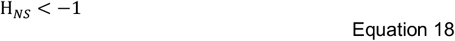

Taken together, it follows that a necessary condition for Equations 1-4 to undergo a Turing instability is that F_NOG_ is a *decreasing* function of [p*S*MAD], i.e., active BMP signaling must inhibit *NOG* transcription. This condition holds regardless of the parameter values chosen in our model, and does not depend on the precise functional form chosen for F_GDF5_ or F_NOG_. Together these results place a rather general constraint on the regulatory logic of the system; this is schematized in Figure 2 in the main text.

#### Extension

In Equations 1-4, we do not take into account the degradation of the complex, since we expect that degradation will be negligible compared to dissociation (i.e., *δ*_COMPLEX_ ≪ *k*_−_). [Molecular half-lives for extracellular ligands are on the order of hours, e.g., (Yelon et al., 2017), whereas off-rates are typically on the order of minutes, e.g., (Nolan et al., 2016)]. Nonetheless, to relax this assumption, we added complex degradation to Equations 1-4 via a term: −*δ*_COMPLEX_[COMPLEX]. When we repeated the linear instability analysis, we could derive the same necessary condition for *b*_1_ < 0. Inspecting the sign of each term contributing to *b*_1_ using MATLAB, we immediately see that the necessary condition can only be satisfied if H_*NS*_ < 0, i.e., F_NOG_ must a *decreasing* function of [p*S*MAD]. Therefore, our main results (Figure 2) still hold if we consider non-negligible rates for complex degradation.

### S3. Model simulations

Whilst we have derived a necessary condition for pattern formation, we do not have a simplified, analytical expression for a sufficient condition. We therefore turned to simulations to explore whether the system is indeed capable of self-organizing Turing patterns.

We began by simulating Equations 1-4 on a static, one-dimensional domain. Initially we neglected digit growth to focus on the intrinsic, pattern-forming ability of the system, and expected that many qualitative features of the patterns (e.g., phase differences between model variables) would be correctly predicted by simulating on a static geometry.

We chose repressive Hill functions to represent the decreasing functions F_GDF5_ and F_NOG_, i.e.,

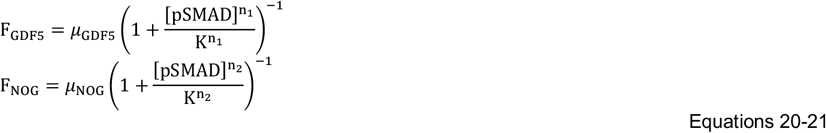

The resulting PDEs are solved using custom MATLAB scripts which are available at https://github.com/twhiscock/bmp_turing_joint_patterning. We use a pseudo-spectral implicit-explicit numerical method to solve the stiff PDEs. Briefly, we discretize the space into N points, which transforms the PDEs into a system of ODEs. At each time step, we combine the (implicit) backward Euler algorithm with the discrete cosine transform (DCT) to efficiently compute the increments associated with diffusion (and assuming reflective boundary conditions). We then use the (explicit) forward Euler algorithm to compute the increments associated with the reaction terms. We use homogeneous initial conditions with low amplitude noise added (using normally distributed random numbers) and explore whether the system can then self-organize into periodic patterns.

To visualize the results, we use line plots and heatmaps that describe the predicted concentrations of the extracellular species (GDF5, NOG and complex) as well as the level of BMP signaling (pSMAD). To compare our *in silico* predictions to *in vivo* mRNA measurements, we assume that the term F_GDF5_ is proportional to *GDF5* mRNA levels, and F_NOG_ to *NOG* mRNA levels (see Fig 3B).

#### Incorporating growth dynamics and cell fate commitment

Having investigated the pattern-forming ability of Equations 1-4, we considered how growth dynamics and cell fate commitment would impact the patterning dynamics. We consider a highly simplified scenario to approximate digit growth, following the approach taken in (Scoones & Hiscock, 2020). Briefly, we assume that the digit both elongates at its distal edge, as well as uniformly stretches along its length. The combination of distal and uniform growth causes the digit length, *L*, to increase over time. In addition, we assume that once cells are above a certain distance, *L*_pattern_, from the digit tip, their concentrations remain fixed in time (i.e., are no longer governed by Equations 1-4) but can still influence patterning in the distal domain due to diffusion. We emphasize that these assumptions are highly unrealistic and fail to account for the complex processes involved in digit outgrowth, morphogenesis, and cell fate determination. Nonetheless, these crude approximations provide us with proof-of-principle simulations demonstrating how growth might impact predicted expression dynamics.

We considered two types of boundary condition at the growing distal tip, either:

##### Reflective-boundary

At each timepoint, we project the concentrations at the digit tip to extend throughout the rest of the domain to enforce reflective boundary conditions (MacDonald et al., 2013).

##### PFR-boundary

Here we assume that, outside of (i.e., distal to) the digit domain, the reaction-diffusion dynamics take a simpler form:

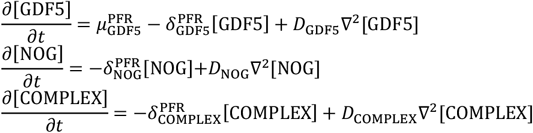

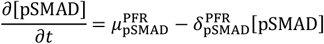

This enforces high pSMAD activity at the distal tip to mimic the PFR. (Note, the above equations are designed to generate a distal pSMAD+ domain as simply as possible, and are not meant to accurately capture the specification of the PFR, which is a more complex process involving other signaling molecules and mechanical cues).

We found that, regardless of which boundary condition we used, the growth dynamics caused patterning to occur exclusively at the distal tip, with new *GDF5* bands being sequentially added as the digit elongates. These spatiotemporal dynamics were observed for a broad range of parameters, although some parameters (e.g., too low/high values of the growth rate or *L*_pattern_) led to patterns that were not observed *in vivo*, such as splitting of phalanges or failure to form repeated patterns, something which is expected from previous studies (Scoones & Hiscock, 2020).

Our choice of boundary condition impacted the predicted expression patterns towards the digit tip. For the *reflective-boundary*, we predicted that *GDF5* expression would initiate at the very tip when pSMAD was low, which would then be followed by initiation of pSMAD at the tip and downregulation of *Gdf5*. In contrast, for the *PFR-boundary*, we see that pSMAD is always high at the tip (mimicking the PFR), and a nascent *GDF5* band first appears proximal to this region, more closely matching the dynamics that we observe *in vivo* (Figure S3C).

#### Model parameters

All model parameters used in this manuscript are provided in the accompanying MATLAB scripts (https://github.com/twhiscock/bmp_turing_joint_patterning). Here we will briefly outline the rationale for our parameter choices.

To explore the pattern forming capability of our model, we simulated Equations 1-4 on a static domain and saw that a wide range of parameters led to periodic patterns. We show some example parameter sets in Figure S2B, which illustrate how different combinations of diffusivities are compatible with patterning.

For simplicity, for the remaining simulations in the manuscript we focused on a single parameter set, although expect the results to be qualitatively similar for other parameter sets. Figure 3B shows the predicted patterns when simulating with these parameters on a static domain. Figure S5B, also on a static domain, takes the same parameter values as Figure 3B, but varies each one individually over a five-fold range (both lower and higher than the reference). We observe that periodic patterns form reliably across a range of each parameter, but that some parameters significantly affect the wavelength of the resulting pattern.

Using the same parameter set, we performed simulations which also incorporated growth dynamics and cell fate commitment. In Figure S3A&C we implement the *PFR-boundary* and predict the sequential addition of new *GDF5* bands at the growing end of the digit. In Figure 4E-G, we mimic experimental perturbations and predict their effect on the *GDF5* pattern (4E: *μ*_p*S*MAD,_ = 0; 4F: *μ*_NOG+_ = 0; 4G: assume extra, localized source of GDF5). In Figure 5B, we explore factors that impact joint number, including variations in growth dynamics (“faster growth”: increase growth rate by 1.333X; “longer duration”: increase patterning duration by 1.333X) and reaction parameters (“shorter wavelength”: increase *δ*_NOG_ by 2X).

## Footnotes

1 Reprinted from Developmental Biology 209(1), E. Storm and D. Kingsley, *GDF5 Coordinates Bone and Joint Formation during Digit Development*, pp. 11-27, Copyright (1999) with permission from Elsevier.

2 From L.J. Brunet, J.A. McMahon, A.P. McMahon and R.M. Harland. *Noggin, cartilage morphogenesis, and joint formation in the mammalian skeleton*. Science 280(5368) pp.1455-1457 (1998). Reprinted with permission from AAAS.

